# EVR: Reconstruction of Bacterial Chromosome 3D Structure Using Error-Vector Resultant Algorithm

**DOI:** 10.1101/401513

**Authors:** Kang-Jian Hua, Bin-Guang Ma

## Abstract

More and more 3C/Hi-C experiments on prokaryotes have been published. However, most of the published modeling tools for chromosome 3D structures are targeting at eukaryotes. How to transform prokaryotic experimental chromosome interaction data into spatial structures is an important task and in great need. We have developed a new reconstruction program for bacterial chromosome 3D structures called EVR that exploits a simple Error-Vector Resultant (EVR) algorithm. This software tool is particularly optimized for the closed-loop structural features of prokaryotic chromosomes. EVR can be used to reconstruct the bacterial 3D chromosome structure based on the contact frequency matrix derived from 3C/Hi-C experimental data quickly and precisely.

## INTRODUCTION

Like eukaryotes, the prokaryotic genomes also fold with regular patterns. The spatial organization plays very important roles in many cellular activities such as transcriptional regulation (1), gene expression (2) and DNA replication (3). Many approaches, *e.g.*, FISH (Fluorescence in situ hybridization) (4) and 3C (chromosome conformation capture) technology (5) have been adopted to study the 3D architecture of prokaryotic chromosome. In FISH experiments, fluorescence is generated by binding a labeled probe to a specific DNA fragment, and the position of target DNA fragment in cell can be visually observed under a microscope (6). However, due to the low throughput and limited resolution of FISH, it is difficult to be applied in the genome-wide study of chromosome structure. The 3C technology and its derivative technologies such as 4C (circular chromosome conformation capture) (7), 5C (3C-carbon copy) (8) and Hi-C (High-throughput/resolution chromosome conformation capture) (9) have significantly promoted the research of three-dimensional genomics in the past decade (1). In 3C technology, the genome is cross-linked by formaldehyde and then broken into fragments which are further ligated by enzymes and sequenced for subsequent analysis (10). In the process of cross-linking, the probability that the fragments close to each other in space are connected is high. This feature is an important clue to study the spatial structure of chromosome (11). In 2011, Umbarger *et al.* used 5C technology to generate a genome-wide DNA interaction map of *Caulobacter crescentus* with fragment size of approximately 13kb (12), and established the 3D structure of chromosome using the integrated modeling platform (IMP) (13,14). Subsequently in 2013, Cagliero *et al*. obtained the chromosome interaction data of *E. coli* (15), but the final interaction map contained a large amount of noise, and it was difficult to use the data to reconstruct structures. In the same year, Le *et al.* produced the full-genome interaction maps of 10kb resolution for *C. crescentus* using Hi-C and built its spatial structure with their own method (16). With this research, they demonstrated the existence of CID (chromosomal interaction domains) boundaries and their roles in gene expression. In 2015, Marbouty *et al.* applied Hi-C to the chromosome research of *Bacillus subtilis* (17) and reconstructed 3D models with the ShRec3D program (18), and then they combined the 3D genome structure with super-resolution microscopy to unravel the role of chromosome 3D folding pattern in the regulation of replication initiation, chromosome organization and DNA segregation. In the meanwhile, Wang *et al.* also studied *B. subtilis* with Hi-C and a large number of chromosome interaction maps were generated at various physiological stages (19,20), and their research revealed the mechanism of SMC condensin complexes in compacting and resolving replicated chromosomes. Recently, Lioy *et al.* succeeded in applying Hi-C in the study of *E. coli* and obtained interaction maps in a high resolution of 5kb, and the 3D chromosome structures were reconstructed with the ShRec3D program (21) to demonstrate the role of nucleoid associated proteins (NAPs) in chromosome organizing.

Years of investigation have yielded many discoveries of bacterial chromosome features, such as cyclic pseudo-nucleation (22), supercoiled regions (23) and plectoneme structures (16), *etc*. With the knowledge of these features, scientists can build the structure of bacterial chromosome by combining experimental data with elaborately developed calculation methods. In 2017, a research work divided *E coli* chromatin into plectoneme-abundant regions and plectoneme-free regions based on ChIP-chip data (24), and built its structure by using multi-scale modeling and Brownian dynamics approaches, resulting in a chromosome model with a resolution of 1NTB (25). More recently, another study constructed a lattice model of *E. coli* chromatin with resolution of 20bp by giving the position and size of the plectoneme fragments (26) using a mesoscale modeling pipeline method (27).

With the application of 3C technology, more and more chromosome interaction maps of prokaryotes emerge and the 3D models of chromosome structure are in great need for the studies of chromosome function and transcriptional regulation. Several software tools have been published for the reconstruction of chromosome 3D structures. These tools can be classified into two categories according to their modeling strategies: modeling based on restraints and modeling based on thermodynamics (28). Restraint-based modeling is a commonly used method to derive 3D models from DNA interaction data. In restraint-based modeling, the DNA interaction data is first converted to the expected space distance, and then heuristic methods are used to optimize the structure under given restraint conditions, such as the gradient ascent method in MOGEN (29), Monte Carlo sampling in TADbit (30), multidimensional scaling algorithm in miniMDS (31), and shortest-path method in ShRec3D (18). In these methods, ShRec3D (18) has been applied to reconstruct the chromosome 3D structures of two model bacteria species (*B. subtilis* and *E. coli*) (17,21), but it is sensitive to resolution and highly depends on data normalization.

In this study, a new algorithm was proposed specially for the chromosome structure reconstruction of prokaryotes, which used error vectors to guide structural optimization and added constraints of a circular genome shape on the implementation of program. This program can better deal with prokaryotic data and has low sensitivity to noise and high calculation speed by fully utilizing the computing power of multi-core CPUs and GPUs. With this algorithm, the 3D structures of the chromosomes of three prokaryotic model organisms (*C. crescentus, B. subtilis* and *E. coli*) were reconstructed and analyzed.

## MATERIALS AND METHODS

### Data source

The sequencing data from 3C/Hi-C experiments can be processed by pipelines such as HiC-Pro (32) or HiCUP (33) to obtain chromosome interaction information. In these pipelines, genomes are divided into equal-length segments and each segment is called a bin. The interaction information of bins is generally presented in the form of a matrix whose element is the number of interactions between two bins. Then the matrix will be normalized to remove some biases such as GC content, mappability, and fragment length, *etc.* (34), resulting in the interaction frequency (IF) matrix, which will be used to construct the spatial structure of chromosome (namely, determine the spatial coordinates of each bin).

We used the published chromosome interaction data of three bacteria species: normalized chromosome interaction frequency matrix (referred to as IF matrix) with 10kb resolution of *Bacillus subtilis* (GEO accession number: GSE68418) (19), normalized IF matrix with 10kb resolution of *Caulobacter crescentus* (GEO accession number: GSE45966) (16), and the raw IF matrix with 5kb resolution of *Escherichia coli* (GEO accession number: GSE107301) (21).

### Algorithm

The EVR algorithm consists of the following steps: (i) transform IF matrix to the expected distance matrix *D* by *Eq.* 1; (ii) an initial conformation is generated by randomly assigning coordinates to DNA bins; (iii) calculate error vectors for each bin; (iv) a new conformation is obtained by moving bins according to the guidance of error vectors; (v) repeat steps (iii) and (iv) until the iteration stop condition is satisfied. The chromosome structure is the final conformation (coordinates of bins) after iteration (**Fig. 1**).

**Fig. 1.**
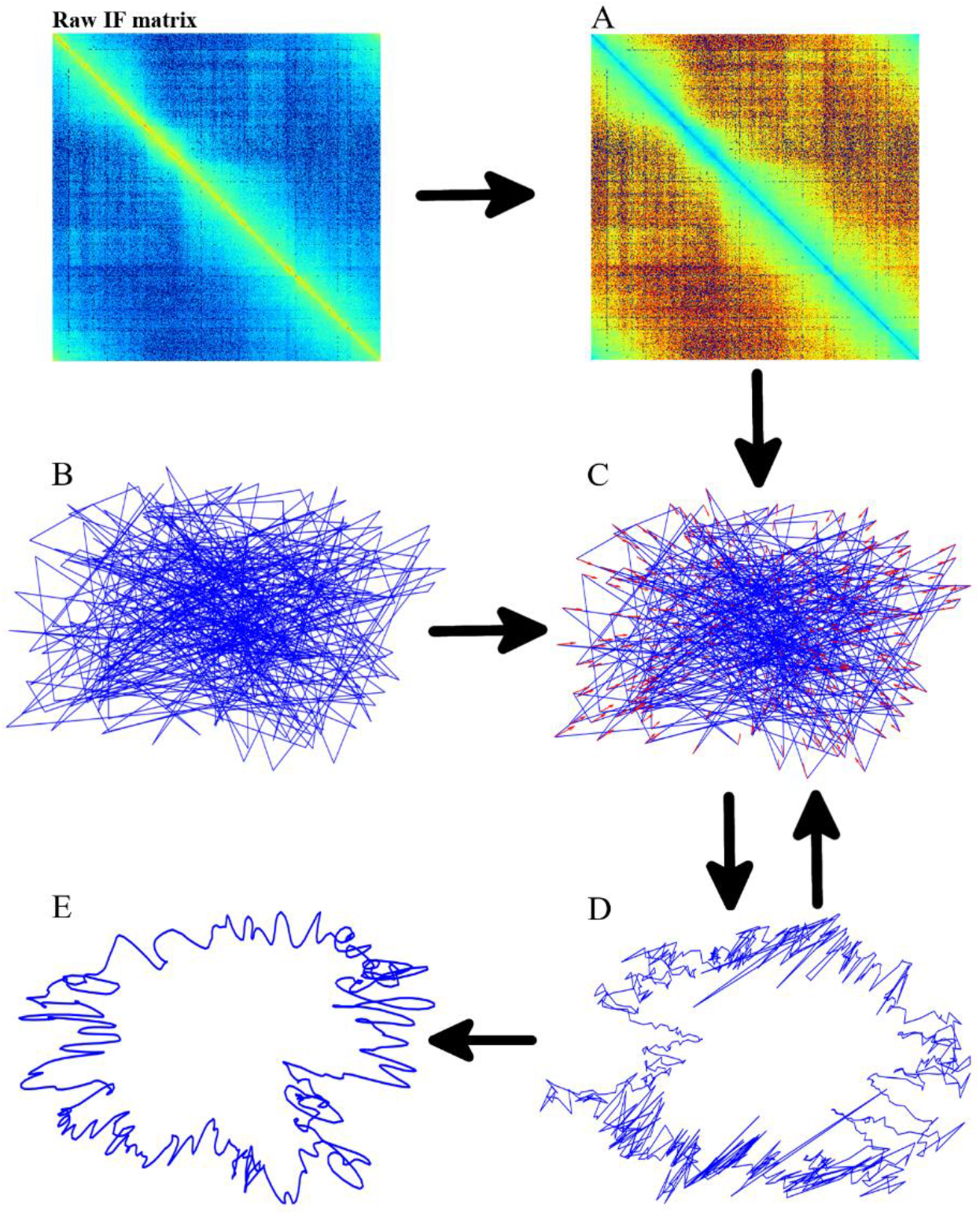
Data flow chart of the EVR algorithm. (A) Expected distance matrix. (B) An initial random conformation. (C) Error vectors on bins: each red arrow is an error vector of a bin. (D) New conformation after moving bins according to the guidance of error vectors. (E) The final structure.

### Generation of the expected distance matrix

In cross-linked chromatin, the closer the spatial distance is, the higher the probability of cross-linking is. Therefore, *Eq.* 1 is usually used to express the relationship between interaction frequency and the expected spatial distance (11). In *Eq.* 1, *F*_*ij*_is the interaction frequency of bin *i* and bin *j*, and *D*_*ij*_is the corresponding expected spatial distance. If *F*_*ij*_ is not equal to 0, a power function is used to convert *F*_*ij*_ into *D*_*ij*_; otherwise, the power function is invalid, and then the distance constraint of adjacent bins will be used to optimize the coordinates. Here, the power exponent *α* is set to 0.5 by reference (21).

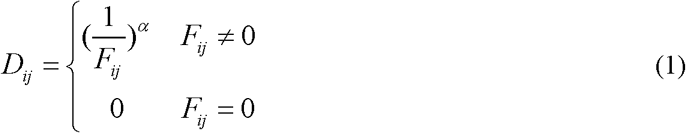

### Error vectors

An initial chromosome conformation is generated by randomly assigning coordinates to all bins. With the distance matrix *D*, error vectors are calculated between every two bins. The error vectors are expressed as in *Eq.* 2 for bins that are not directly adjacent on the linear genome. Here,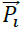 and 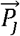 are the coordinates of bin *i* and bin *j* in vector form, respectively, and *D*_*ij*_ (*Eq.* 1) is the expected distance between the two bins.

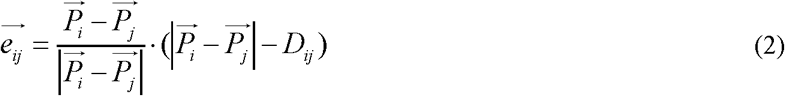

For bins that are directly adjacent on the linear genome, error vectors are defined as in *Eq.* 3, where *D*_*max*_ and *D*_*min*_ are the allowed maximum and minimum distances between any two adjacent bins, respectively. In a conformation, *D*_*max*_ and *D*_*min*_ will be used to calculate the error vector between two adjacent bins if the distance of the two bins is less than *D*_*min*_ or greater than *D*_*max*_; otherwise, the expected distance *D*_*ij*_ is used. To cope with the cyclic feature of bacterial chromatin, the first bin and the last bin are set as adjacent bins.

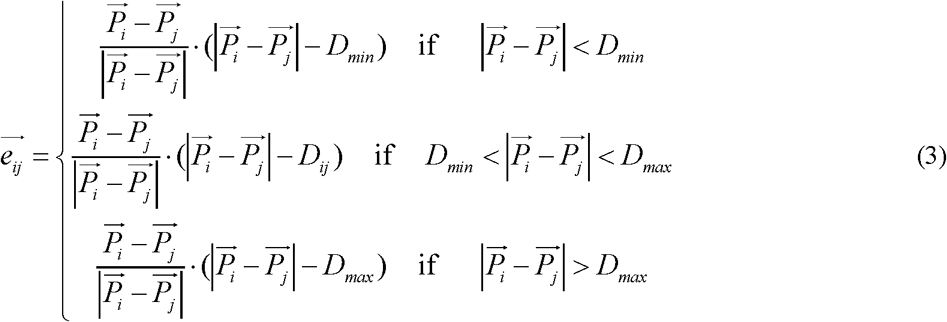

For each bin, all its error vectors are summed up with certain weights to a resultant vector (EVR). As shown in *Eq.* 4,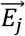is the EVR for bin *j* which indicates the direction and distance that bin *j* should move in the next iteration step (**Fig. 2A**) and a weight *w*_*ij*_ determines the step size. When *w*_*ij*_ is large, the length of EVR is large, and the structure changes greatly compared with the previous one after one iteration step, and the optimization speed is fast, but the result may not converge. Conversely, when *w*_*ij*_ is small, the structural change in each iteration step is small, and the optimization speed is slow, but it is easier to converge. In our algorithm, *w*_*ij*_ = 1/*N*, where *N* is the number of bins.

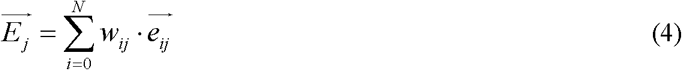

**Fig. 2.**
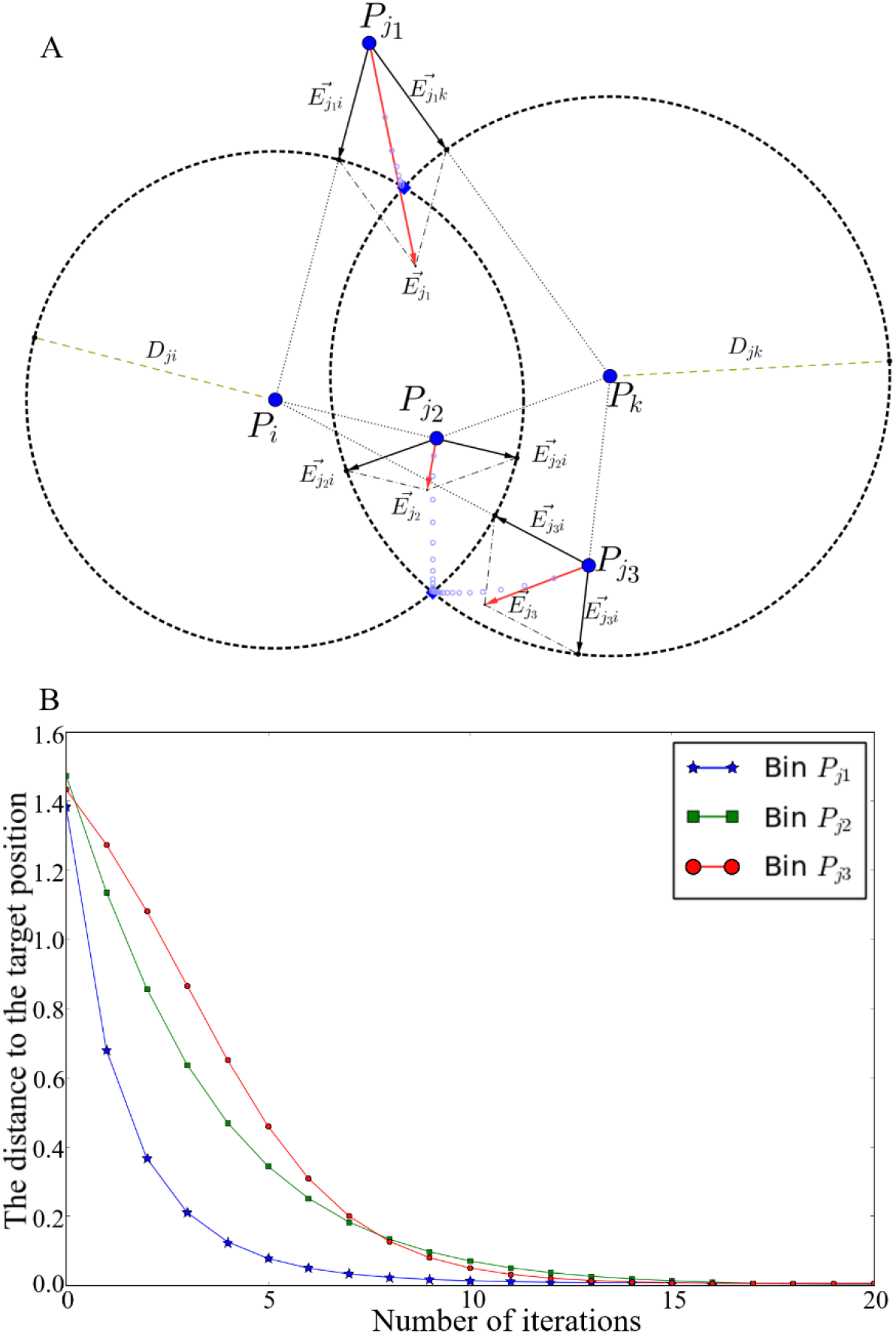
EVR calculation. (A) Illustration of error-vectors and error-vector resultants. *P*_*j*1_,*P*_*j*2_ and *P*_*j*3_ are 3 bins whose positions (coordinates) need to be adjusted; *P*_*i*_ and *P*_*k*_ are 2 bins whose positions are fixed; *D*_*ij*_ and *D*_*jk*_ are the expected distances between these 3 bins to bins *P*_i_ and *P*_*k*_; the intersection points of the two dotted circles (centered at *P* _*i*_and *P*_*k*_ with radii *D*_*ij*_ and *D*_*jk*_, respectively) are the target positions of the 3 bins *P*_*j*2_, *P*_*j*1_ and *P*_*j* 3_; the closest intersection point is chosen as the target position for each of the 3 bins. (B) The distances between the 3 bins *P*_*j* 1_, *P*_*j*2_, *P*_*j*_ 3 to their target positions decrease during iteration.

An example of EVR calculation is shown in **Fig. 2A**, in which black arrows are error vectors that can be calculated according to *Eq.* 2 and *Eq.* 3 and red arrows are the resultant vectors calculated according to parallelogram law; these resultant vectors can guide the position optimization of bins.

### Iterative optimization

Starting with an initial random conformation, error vectors and resultant vectors can be calculated based on the expected distance matrix for all bins. The resultant vector of each bin indicates the direction and distance according to which the bin should move to change its position towards the target position. The chromosome structure is optimized by an iteration process during which all bins change their positions (coordinates) guided by their resultant vectors, resulting in the minimization of the sum of EVR lengths (*Eq.* 5). For example, the motion trajectories of the 3 bins *P*_*j* 1_, *P*_*j* 2_, *P*_*j* 3_ are shown as light blue circles (**Fig. 2A**) and the distances of the 3 bins to their target positions decrease quickly during iteration (**Fig. 2B**).

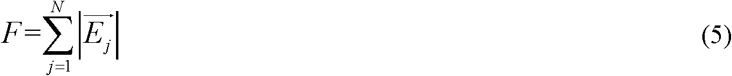

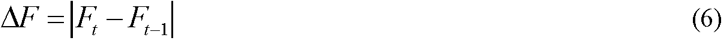

In the iterative process, the value of F becomes smaller and smaller to approach a stable value. As shown in *Eq.* 6,ΔF is defined as the difference between two successive F values of iterative steps *t* and *t*-1. When a level of ΔF < 1E-5 or the given maximum number of steps is reached, the iteration stops and an optimized conformation is obtained which is the final structure of a chromosome.

### Implementation

The main part of EVR program is implemented in Python with Numpy arrays for storing data. Two acceleration schemes are adopted in the iterative optimization part: Cython and OpenCL. In Cython scheme, the core code is written in Cython-C language and compiled into a dynamic library that can be accelerated by OpenMP. The Cython scheme can make full use of the computing power of multi-core CPUs. In OpenCL scheme, the core code is written in OpenCL-C language, and it can be compiled and invoked directly by the PyOpenCL library functions. The OpenCL scheme can use not only multi-core CPUs but also GPUs and other devices that support OpenCL to accelerate computing.

## RESULTS

First, we compared our EVR program with some published software tools for chromosome reconstruction, including: miniMDS (31), MOGEN (29) and ShRec3D (18). All these software tools ran on multi-core CPUs with default parameter setting. The comparisons involve the speed, accuracy and sensitivity to noise level of calculations. Subsequently, our EVR program was applied to the chromosome structure reconstruction of 3 model species of prokaryotes with published 3C/Hi-C data. The obtained chromosome structures were analyzed in a biological background.

### EVR is fast

Among the compared software tools, miniMDS is written in Python and accelerated using the pymp library; MOGEN is written in java and can use multi-core CPUs for accelerating calculation; ShRec3D is written in MATLAB, and the program itself is not parallelized, but the new version of MATLAB supports automatic parallelization. Following convention (11), standard structures were generated for algorithm assessment. Here, toroidal spiral curves were generated and cut into fragments (bins) with bin numbers from 100 to 5000, and the IF matrixes of these structures were calculated to simulate a prokaryotic circular chromosome. Then, the above software tools were used to reconstruct the standard structures with these IF matrixes. See **Supplementary Information** for more details. The time efficiency of structure reconstruction is shown in **Fig. 3.**

**Fig. 3.**
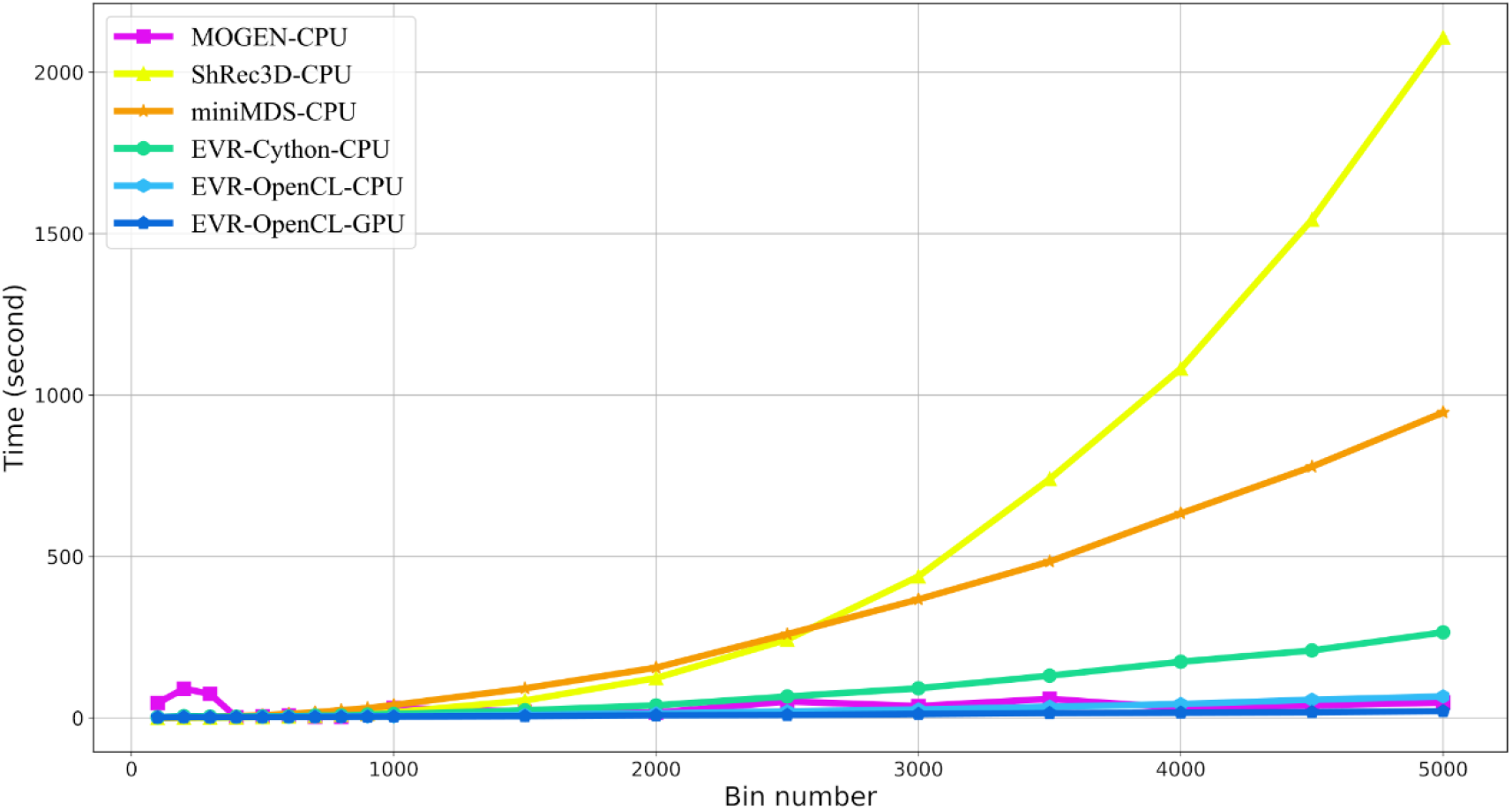
The required time for standard structure reconstruction with different bin numbers.

Testing platform is Ubuntu 16.04 (64 bit) with Intel Core i5-2400 CPU@3.10 GHz and 16GB DDR3-1600 memory. The GPU platform is Geforce GTX 1050Ti. The comparison results show that EVR running on GPU with OpenCL used the least time, followed by EVR running on CPUs with OpenCL and MOGEN; meanwhile, their computation time is not sensitive to the bin numbers within a reasonable range. The running speed of EVR with Cython is slightly slower but still relatively fast. The computation time of miniMDS and ShRec3D shows a rapid rise with the increase of bin numbers.

### EVR is robust

To assess robustness, different levels of noise were added to the IF matrix generated from the structures of toroidal spiral curves, and then the noisy data were used in the 3D structure reconstruction by using the above four software tools. Noises were added to the IF matrix by randomly selecting values from the range of −0.5 × *P* × *IF*_*max*_ to 0.5 × *P*× *IF*_*max*_; here, *P* is noise level and *IF*_*max*_ is the maximum value in IF matrix. The resulting structure reconstructed from the noisy data was compared to the original noise-free structure and the RMSD (root-mean-square deviation) value was calculated. See **Supplementary Information** for more details. At each noise level, the above steps of adding noise, reconstructing structures, and comparing structures were repeated 100 times. The average RMSD values obtained are plotted against noise level as **Fig. 4**.

**Fig. 4.**
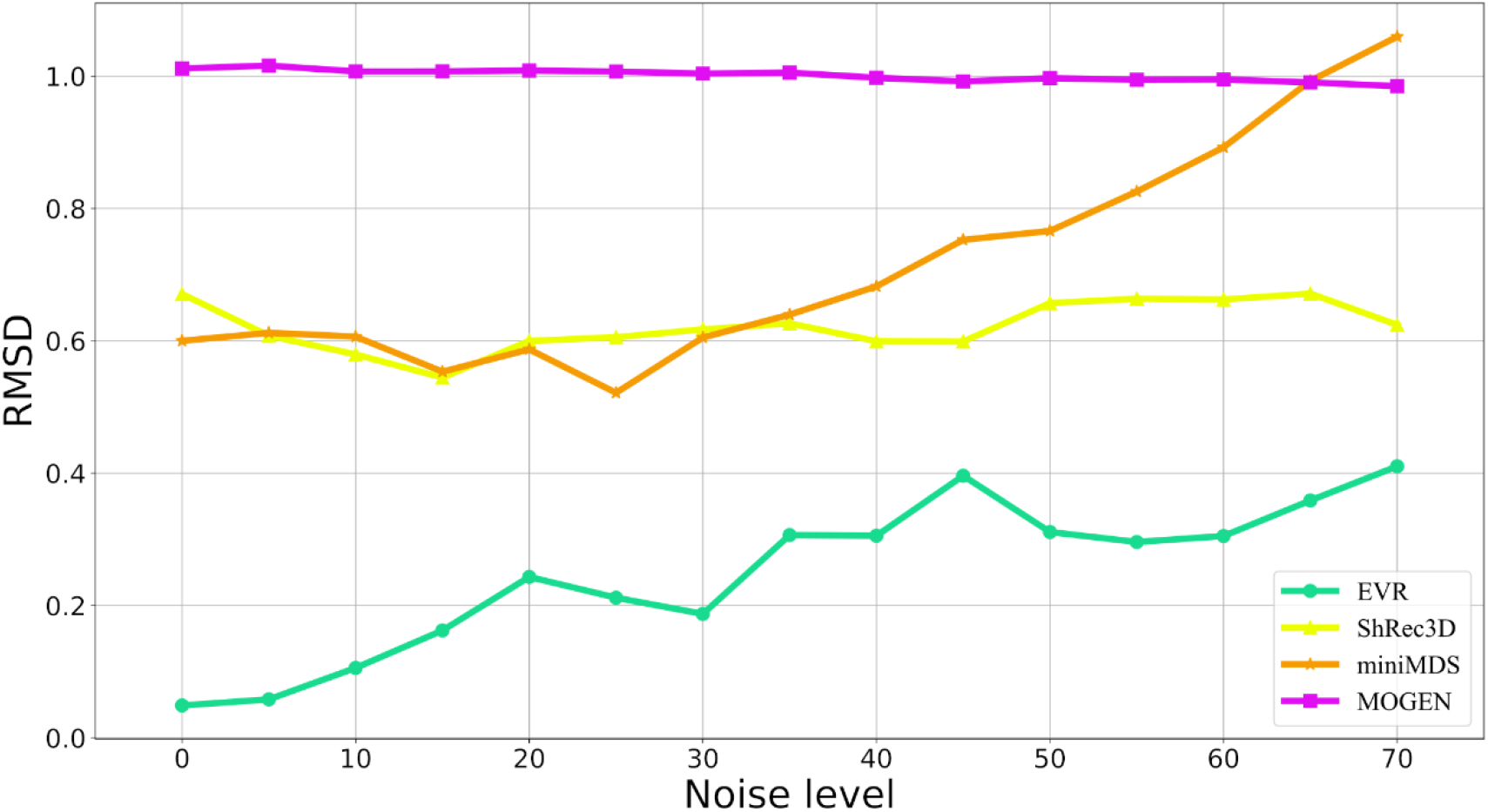
Comparison between reconstructed structures (from noisy data) and original structures using four software tools. Both the absolute RMSD value and its trend with the increase of noise level should be considered.

**Fig. 5.**
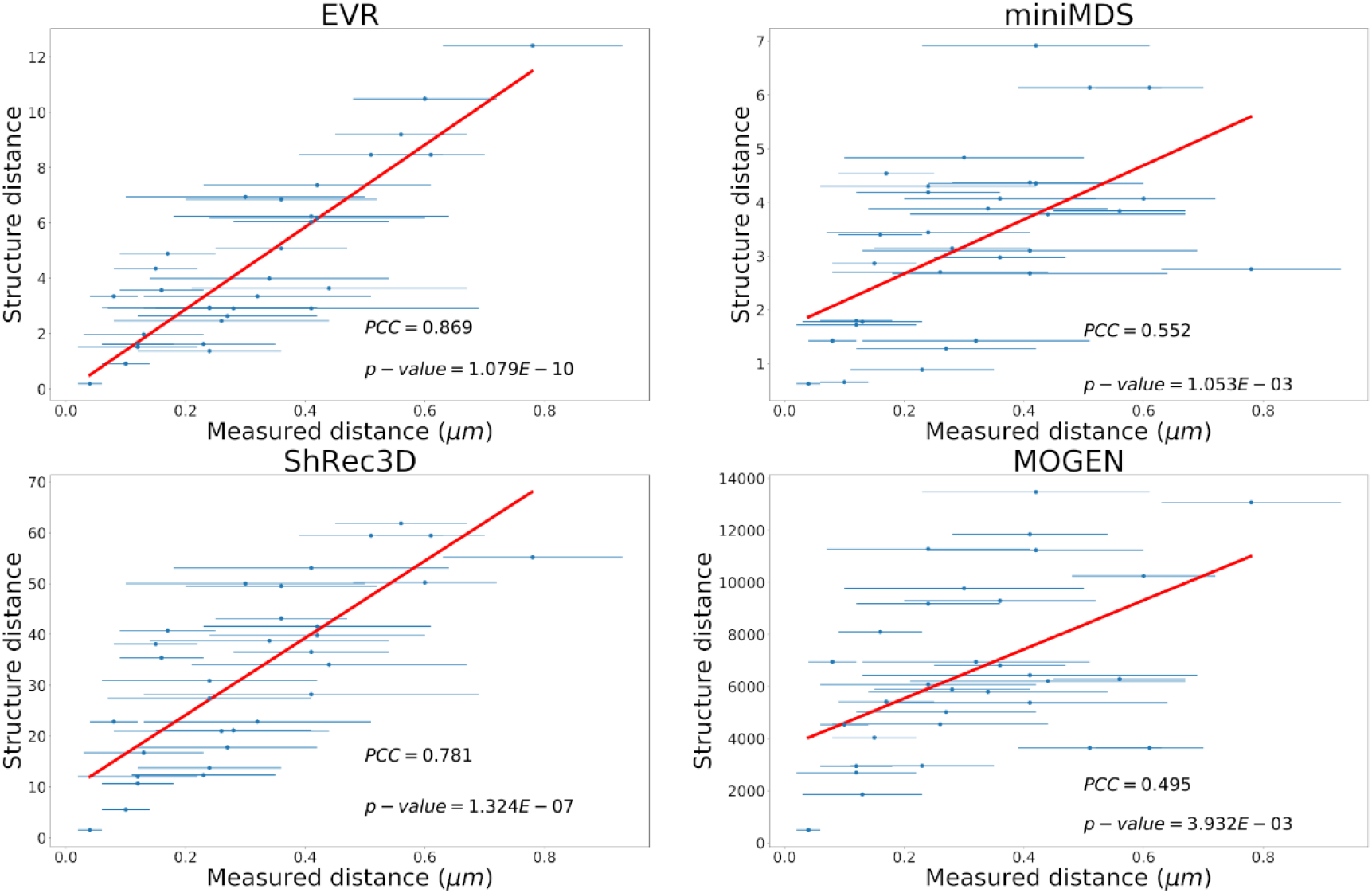
Correlation between the structure-based distance and the experimentally measured distance. The chromosome structures were reconstructed by using the four software tools based on recently published 3C data of ***E. coli*** (21) and the experimentally measured distances were compiled from literature (35). See **Supplementary Information** (**Table S1**) for more details.

Since the IF matrix does not provide structural scale information, the scales of the structures obtained by different software tools may be not the same. When comparing structures, in order to ensure scale consistency, all the structures are scaled and compared with the original structure, and the structural alignment with the smallest RMSD value is selected for comparison. With this comparison strategy, even if two structures are very different, their RMSD values will be less than around 1.0. As shown in **Fig. 4**, our EVR algorithm always has the smallest RMSD values at different levels of noise, meaning that the structures reconstructed based on the noisy data are always very similar to the original structures based on the noise-free data, i.e., our EVR algorithm is robust to noise.

### EVR is accurate

To assess accuracy, the reconstructed chromosome structures of *E. coli* by using the four software tools were compared with the published experimental fluorescence microscopy data (**Table S1**) (35). The *E. coli* chromosome structure reconstruction was based on the recently published IF matrixes (21). By mapping the fluorescence marker sites onto the reconstructed structures (**Fig. S4**), the 3D coordinates of these sites could be determined and the spatial distances between them were calculated and correlated with the experimentally measured distances in fluorescence microscopy. Results show that there are high correlations between the structure-based distances and the experimental distances for the EVR (*PCC* = 0.869, *p* - *value* = 1.08 *E* - 10) and ShRec3D (*PCC*= 0.781, *p* – *value* = 1.32 *E* - 7) software tools; The correlations for miniMDS (*PCC* = 0.552, *p*- *value* = 1.05*E* - 3) and MOGEN (*PCC* = 0.495, *p* - *value* = 3.93 *EHH* - 3) are relatively lower but still significant, possibly because they are developed for eukaryotes, not suitable or optimized for prokaryotes. The significant and high correlation between the experimentally measured and structure-based distances indicate that the reconstructed structure is close to the real structure and that our EVR algorithm is accurate.

### Application of EVR to prokaryotes

Our EVR algorithm was applied in the reconstruction of 3D chromosome structures of three prokaryotic model species *Escherichia coli* (MG1655), *Caulobacter crescentus* (CB15) and *Bacillus subtilis* (PY79) based on 3C data, and the obtained structures were divided into 4 macrodomains (1,36): origin and terminus of replication, left and right chromosomal arms (**Fig. 6**).

**Fig. 6.**
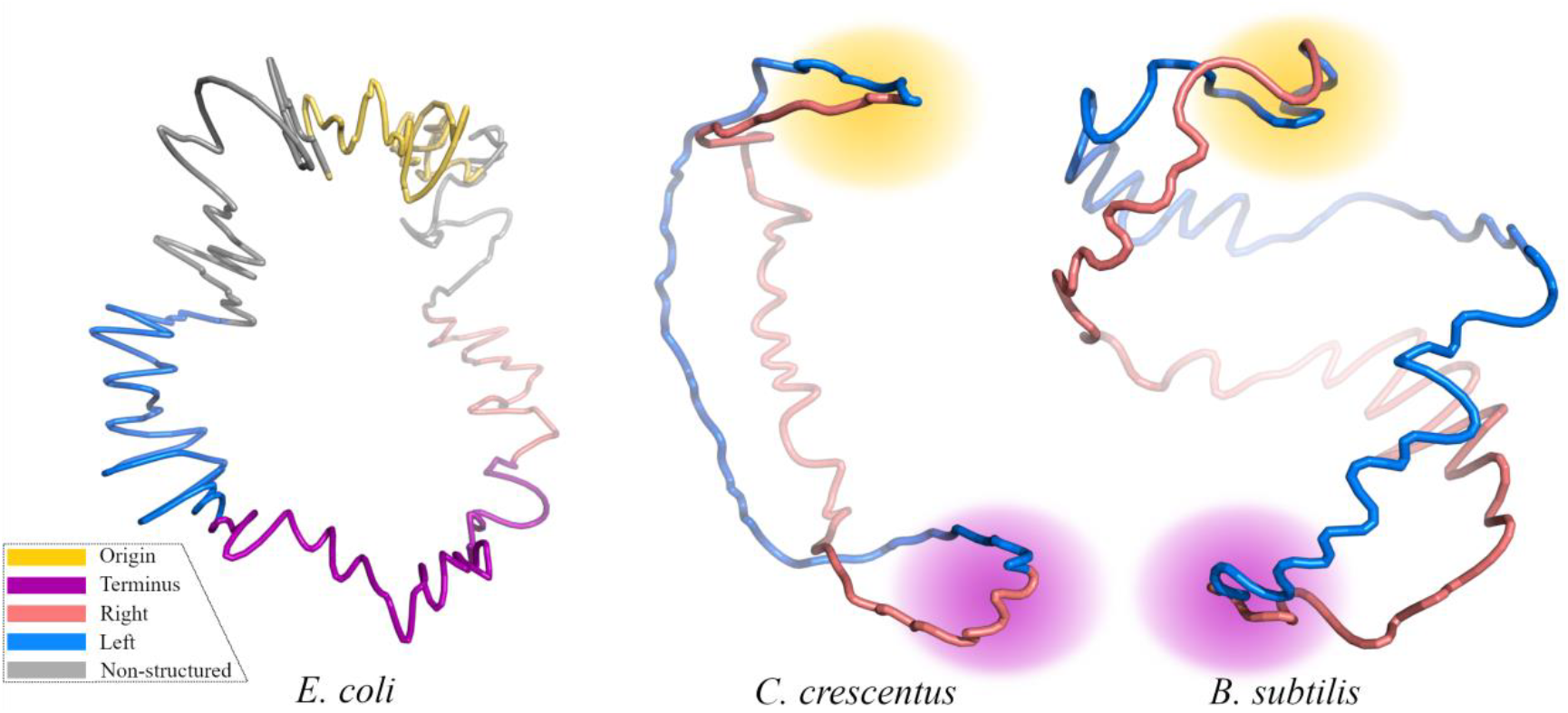
Schematic representation of the reconstructed 3D chromosome structures of three model prokaryotes. Macrodomains are colored as indicated by legend. The macrodomains of *E. coli* are accurately determined according to previous publication (36), while those of *C. crescentus* and *B. subtilis* are roughly shown with halos.

After transforming the interaction matrix into a spatial structure, we can visually examine the structure from a global perspective and observe features that are difficult to find in the interaction matrix. For example, Fis is a very important nucleoid-associated protein in *E. coli*, and it plays a variety of roles in regulating DNA transactions and modulating DNA topology (37). By comparing the interaction matrixes of wild-type and *fis*-deficient (Δ*fis*) *E. coli*, it can be seen that the interaction intensity is significantly reduced at the terminus region but conserved at other sites (21). While converting these two IF matrixes into structures by using our EVR program and comparing them, the overall effect of Fis protein on the 3D structure is even more obvious (**Fig. 7**): compared to the wt type, there is a clear angle in the terminus region of the Δ*fis* type, although other regions are substantially overlapping. This angle cannot be found in the interaction map. Why the lack of *fis* causes the bending of the terminus region and what biological effects this bending will cause remain to be further investigated.

**Fig. 7.**
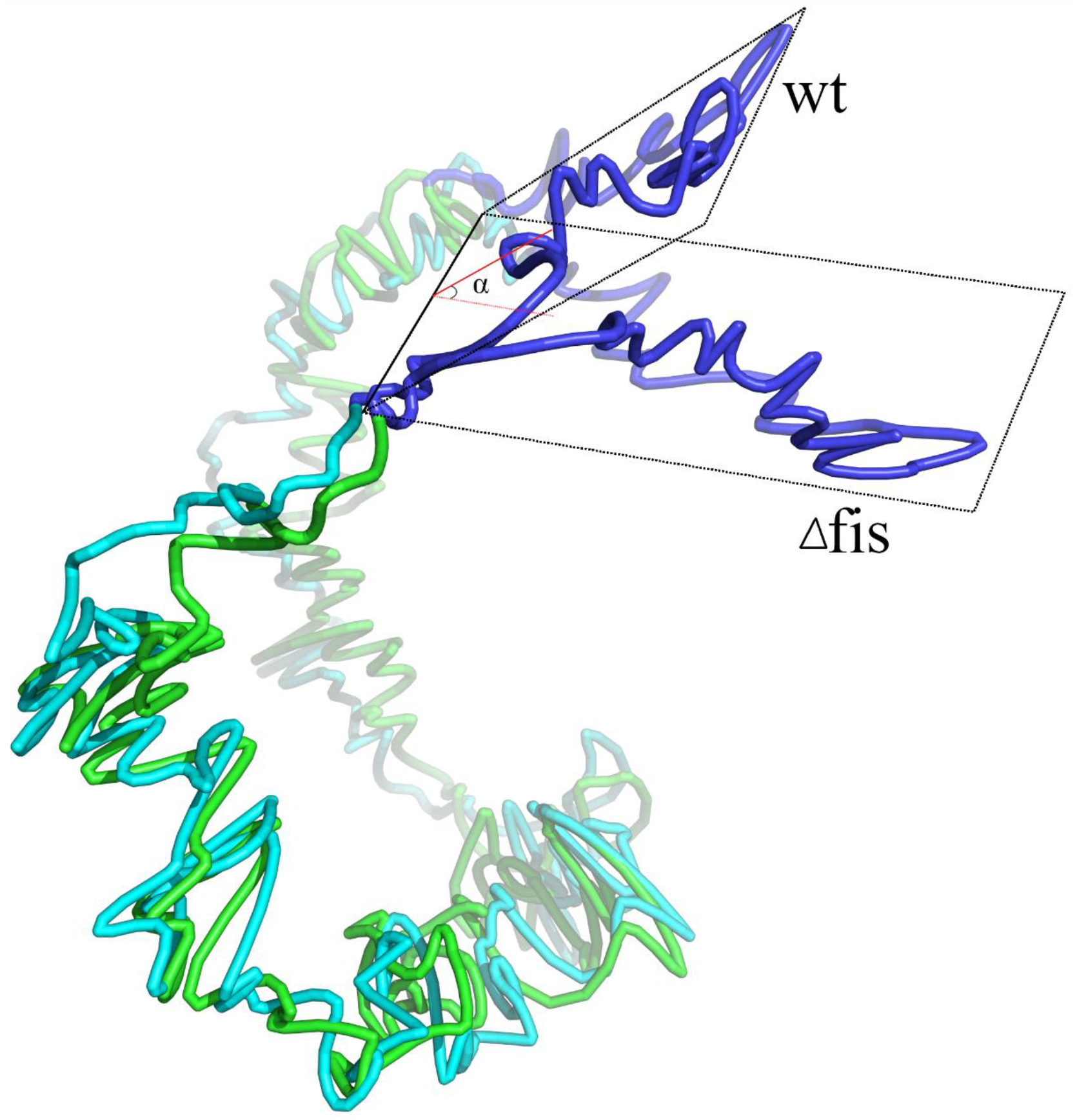
Comparison of chromosome structures of wild-type and Δ*fis* type *E. coli*. Green and cyan tubes represent the chromosomes of wild-type and Δ*fis* strains, respectively, and the blue parts are the terminus region. There is a clear separation between the two structures in terminus regions, which cannot be easily found without accurate 3D structures.

## DISCUSSION

We have developed a software tool called EVR for the reconstruction of 3D chromosome structures from the DNA interaction data measured by 3C/Hi-C experiments. This tool added constraints related to a circular shape in the reconstruction process and thus it is particularly suitable for the reconstruction of chromosome structures of prokaryotes. Assessed based on standard structures, our EVR program is very fast and robust compared with other software tools, and the reconstructed structure is accurate evaluated based on the fluorescence labeling data. Our EVR program was used in the reconstruction of the 3D chromosome structures of three model prokaryotic species with published DNA interaction data. The obtained structures provided consistent understanding on the 3D genomes of these species.

Limited by the resolution of 3C/Hi-C experiment data (38,39), our EVR program as well as other programs can only establish the backbone structure of the prokaryotic chromosomes at domain levels. Combined with other modeling approaches such as the thermodynamics-based methods used in a recent paper (25), more reliable and higher resolution chromosome models may be built to assist the fundamental research on the 3D genomics of prokaryotes. Another noticeable aspect is about the storage file format of the reconstructed chromosome structures. Currently, since the resulting structures contain only the spatial coordinate information of bins, they are usually saved in a simple PDB-file-like format such as the modified PDB file format in (29) or“xyz” format in (18). With the development of 3D genomics, more specific format may be explored to accommodate more information such as domain partition, gene annotation and levels of transcription *etc.* in order to be readily integrated into a systems biology context.

## DATA AVAILABILITY

The source code and data of the program are available at https://github.com/HakimHua/EVR.

**SUPPLEMENTARY INFORMATION**

## ACKNOWLEDGEMENT

We’d like to thank Yang Yi for his help in polishing the manuscript.

